# Fully unsupervised deep mode of action learning for phenotyping high-content cellular images

**DOI:** 10.1101/2020.07.22.215459

**Authors:** Rens Janssens, Xian Zhang, Audrey Kauffmann, Antoine de Weck, Eric Y. Durand

**Author notes:** Authors contributed equally (Antoine de Weck, Eric Y. Durand).

## Abstract

The identification and discovery of phenotypes from high content screening (HCS) images is a challenging task. Earlier works use image analysis pipelines to extract biological features, supervised training methods or generate features with neural networks pretrained on non-cellular images. We introduce a novel fully unsupervised deep learning algorithm to cluster cellular images with similar Mode-of-Action together using only the images’ pixel intensity values as input. The method outperforms existing approaches on the labelled subset of the BBBC021 dataset and achieves an accuracy of 97.09% for correctly classifying the Mode-of-Action (MOA) by nearest neighbors matching. One unique aspect of the approach is that it is able to perform training on the entire unannotated dataset, to correctly cluster similar treatments beyond the annotated subset of the dataset and can be used for novel MOA discovery.

## 1 Introduction

High Content Screening (HCS) (Götte *et al*., 2010; Caicedo *et al*., 2017; Buchser *et al*., 2004) is a phenotypic screening method leveraging high-throughput microscopy imaging to explore and identify perturbations, here low molecular weight compounds, with phenotype altering effects on cells. Experiments are performed on plates with multiple cell-containing wells. The cells are stained with different fluorescent probes highlighting distinct cellular components. Each well is treated with compounds at various concentrations. The goal is to perform quantitative analysis of the resulting images to uncover what happens in the cells upon treatment. In particular this can yield a better understanding of the mechanism by which cells are affected by various treatments. This is called the Mode-of-Action or sometimes the Mechanism-of-Action (MOA). HCS is a high-throughput method that can generate up to millions of images per experiment, therefore requiring extensive computational analysis.

Several methods have been developed to tackle this problem. For example, Ljosa *et al*., (2013) extract cell embeddings with the Cellprofiler software (Carpenter *et al*., 2006) to predict the MOA based on nearest neighbors’ annotation and Kraus *et al*., (2016) uses a supervised convolutional multiple instance learning for microscopy image classification and segmentation. Ando *et al*., (2017) follow a similar idea but use a pretrained deep metric network to extract features from individual cells segmented from larger images. The work of Godinez *et al*., (2017) contains a multi-scale neural network that makes the extraction of cell regions redundant as the neural network picks up local cell features as well as global structures, such as cell density, from the full image. The neural network is trained in a supervised setup on an annotated subset by using known MOAs as labels and applied on the full dataset. In a real-world scenario, we often do not know the MOA, or only have partial understanding of the MOA of treatments, thus limiting the application of the method to well annotated datasets only. This common lack of MoA annotation is another strong limitation of approaches based on a priori knowledge and annotation. In more recent papers, image metadata are used as pseudo-label to train the model in a self-supervised manner (Godinez *et al*., 2018; Spiegel *et al*., 2019), with promising results. Such metadata typically comprise compound identifier, dose and treatment duration. One concern is that the method considers similar treatments (e.g. small dose variation of the same compound) the same as completely different ones (e.g. different MOA). In addition, batch effects could cause images with identical metadata to look different. All the methods above, besides Ljosa *et al*., (2013), use deep neural networks trained either in a supervised (Kraus *et al*., 2016; Godinez *et al*., 2017), self-supervised setup (Godinez *et al*., 2018; Spiegel *et al*., 2019) on an annotated subset of the data or make use of a neural network pre-trained on non-cellular images (Ando *et al*., 2017; Tabak *et al*., 2019). Using a neural network trained on non-cellular images yields the risk of domain-specific features being lost in the process of capturing features on data not related to HCS. In parallel, Caron *et al*., (2019) developed an unsupervised method, applied to non-HCS data, achieving results close to supervised ones by using a standard clustering algorithm to create pseudo-labels during training.

Inspired by Caron *et al*., (2019), Ando *et al*., (2017) and Godinez *et al*., (2018), we developed a fully unsupervised neural network for clustering similar phenotypes together, solely based on the intensity values of the cellular images. Based on the findings of Godinez *et al*., (2017), we make the extraction of cellular candidates redundant by using an updated multi-scale neural network. The backend is built on top of the deep clustering framework from Caron *et al*., (2019). Additionally, we show that combining two batch correction methods, Typical Variation Normalization (TVN) (Ando *et al*., 2017) and Combat (Johnson *et al*., 2007), during training, significantly improves the results and creates more representative embeddings. We call our method UMM Discovery (<u>U</u>nsupervised <u>M</u>ulti-scale <u>M</u>ode-of-action Discovery) and show that we obtain state-of-the-art accuracy in BBBC021. To evaluate the proposed method on a real-world scenario, we trained it, unlike other methods, on the entire dataset (without MOA labels) and show that it can be used for novel MOA discovery.

## 2 Methods

### 2.1 Our Approach: Unsupervised Multi-Scale Mode-of-Action Discovery (UMM Discovery)

We developed a fully unsupervised deep learning method to cluster cellular images with similar MOA together. We did so by applying a deep clustering framework developed by Caron *et al*., (2019), called DeepCluster, on cellular imaging. Following Godinez *et al*., (2018) findings, we decided to use an updated version of the Deep Neural Network (DNN) architecture, called Multi-Scale-Net. Together, these two methods create a fully unsupervised algorithm, which we call UMM Discovery, that allows us to cluster similar images together. **Fig. 1** shows an illustration of our combined approach. The algorithm uses no prior knowledge of treatment, beyond their respective identifier and relative concentration. The DNN input consists solely of unannotated images and it outputs one 64-feature vector per image.

**Fig. 1:**
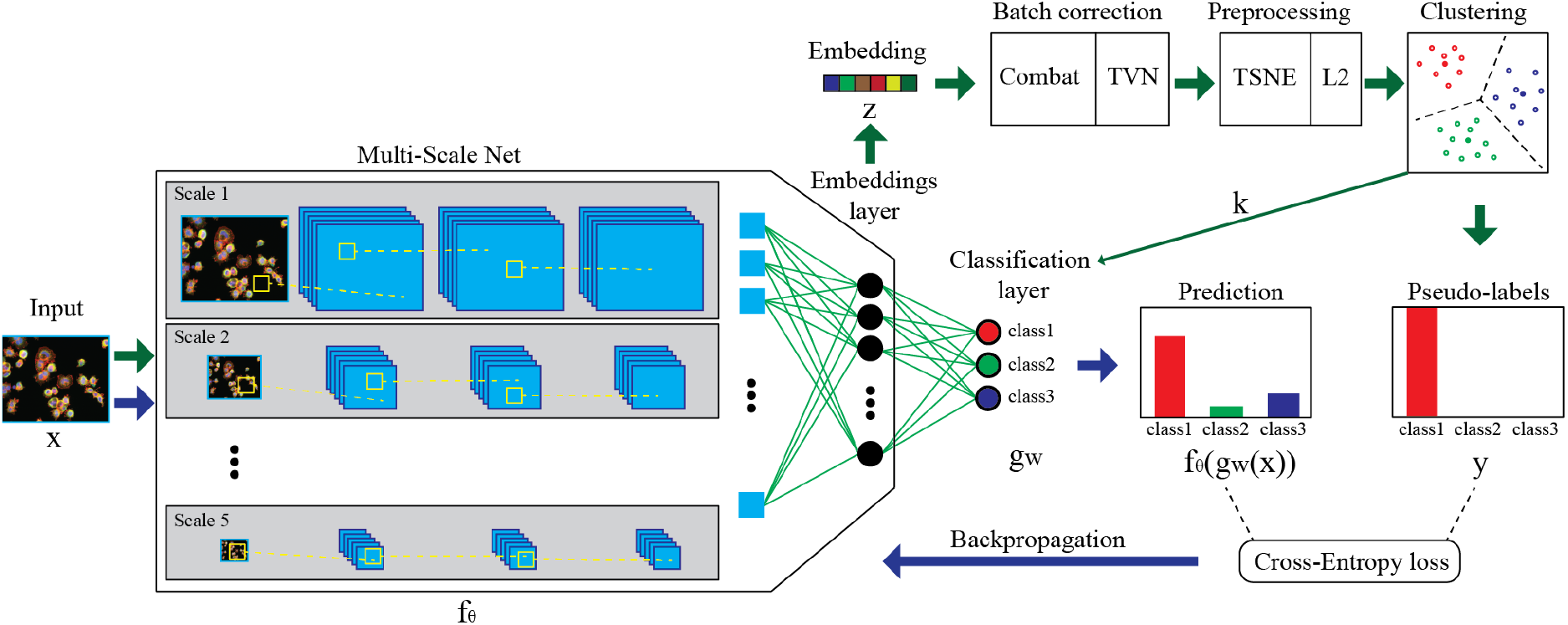
**Overview of the UMM Discovery architecture**, combining an adjusted Multi-Scale NN and the DeepCluster backend. Training of the NN is done in two steps. **Step 1) Green arrows**: Images are fed through the NN where features are extracted from different scales to create one representing embedding for each image. The images are batch corrected with Combat and TVN and preprocessed by TSNE and L2 normalization. The normalized embeddings are clustered with k-means. The cluster assignments from k-means are the new pseudo-labels and the number of clusters will become the size of the classification layer. **Step 2) Blue arrows**: After creation of the pseudo-labels, the training process of the Multi-Scale network starts, where images are forwarded through the NN and the last classification layer to get a prediction for each image. The weights of the Multi-Scale NN are optimized by taking the cross-entropy loss function between the prediction and the pseudo-label and backpropagating the loss through the NN.

#### 2.1.1 Deep Clustering

We built upon the deep clustering framework developed by Caron et al. (2019). The deep clustering framework consists of (i) a neural network for feature extraction, defined as *f_θ_*: *X* → ℝ^*D*^, which takes as input a cellular image *x* ∈ *X* and maps it to a feature embedding *z* of dimension *D* and (ii) a standard clustering algorithm for generating a pseudo-label *y* for each *x*. The training of the parameters *θ* can be summarized as follows. All images *x* ∈ *X* are fed through the neural network to create representative embeddings *z* = *f_θ_*(*x*) ∈ ℝ^*D*^. A dimensionality reduction algorithm is applied to remove dimensions with low variance. Next, a clustering on the embeddings *Z* is performed with a standard clustering algorithm such as k-means to assign each embedding z to one of *K* clusters. The resulting cluster assignment is used as pseudolabel *y* and a classification layer *g_w_* with parameters *w* is added to the end of the neural network, with *K* units representing the *K* clusters, to predict the labels *y* from the feature embeddings *z*. We now have a supervised learning algorithm and can train the neural network with the cluster assignments as pseudo-labels by optimizing:

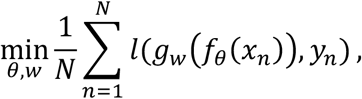

where *l* is the multinomial logistic loss. This process is repeated for each epoch.

#### 2.1.2 Multi-Scale Network

Next, we implemented the multi-scale neural network architecture from Godinez *et al*., (2017) and modified it by adding residual layers, dilation and a collapse layer. The updated version of the Multi-scale network is shown in **Supplementary Figure 1.** The method takes preprocessed images as input and performs a parallel multi-scale analysis over a large number of scales, allowing it to detect local features as well as population changes. This makes it possible to capture the phenotype over a large spatial region. Therefore, no extraction of cellular candidates is necessary.

Given a preprocessed image with spatial dimensions *w* × *h*, the neural network considers downscaled images with dimensions (*w/s* × *h/s*). The neural network contains five multi-scale blocks which reduce the spatial dimension by *s* = 1,2,4,8,16. The multi-scale block consists of three convolutional layers, the first one with a kernel size of 3 and a dilation size of 1, the second one with a kernel size of 3 and a dilation size of 2 and the last one with a kernel size 1. This 1×1 convolutional layer can be seen as a fully connected layer between the feature maps. It is followed by a max pooling operation. Unlike the original multi-scale network from Godinez et al. (2017), the network consists of residual layers. Instead of only considering a downscaled version of the original image, the downscaled image is concatenated with the pooled kernel maps of the last convolution layer and go through an additional 1×1 convolutional layer. After the last multi-scale block, the spatial size is 16×20 pixel with 128 feature maps. A collapse layer is applied, where parallel adaptive average pooling and maximum pooling with the kernel size of the input spatial dimensions. These two layers are afterwards factorized by the trained weight factor *w\** and summed together as follows:

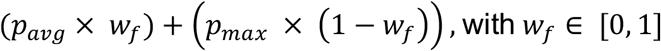

Next, another 1×1 convolutional layer, as well as a fully connected layer with 64 units, are applied. The fully connected layer is followed by a ReLU operation during training. During the embedding generation phase, this ReLU is removed. Batch normalization followed by a ReLU layer is applied after each convolutional layer throughout the whole neural network. With this setup, the neural network takes the images as input and outputs embeddings. The neural network consists of 106’961 trainable parameters, which is low in comparison to other state-of-the-art neural network architectures, for example ResNet-50 with over 25M parameters.

#### 2.1.3 Clustering Algorithms

To generate pseudo-labels, we tested four different clustering algorithms on the embeddings. The first one is the standard clustering framework k-means. The most important parameter for this clustering algorithm is the number of clusters also sometimes called seeds. In the baseline setup the number of clusters is set *a priori*. However, in most real-world scenarios, we do not know the exact number of clusters, therefore we set this parameter to the number of treatments since identical treatments should in theory cluster together.

The true number of clusters, in this case, the number of MOA is unknown. Therefore, two more dynamic clustering algorithms were tested as well. Namely, the Power Iteration Clustering (PIC), an updated version of spectral clustering, and the densitybased method HDBScan (McInnes *et al*., 2017), were tested as clustering algorithms. As noted, the benefit of these two clustering algorithms is that they do not require the number of clusters to be specified. PIC is very fast on large datasets and can provide similar results as spectral clustering. For PIC and HDBScan, we performed hyperparameter optimization to find their best parameters. We also implemented an adaptive version of K-means that reduces the number of clusters linearly during training, starting from double the number of treatments and reducing it to the number of compounds.

#### 2.1.4 Batch Correction

Batch effects arise from uncontrollable variations in biological experiments due to different factors such as slightly different concentrations for same treatments, location of a certain well on the plate, order of the plate within a batch which could result in different rate of evaporation (Tabak *et al*., 2019). To correct for this nuisance variation, we tested two batch correction methods, typical variation normalization (TVN) (Ando *et al*., 2017) and Combat (Johnson *et al*., 2007). Both methods require additional input information. The TVN method needs the embeddings corresponding to the control wells (untreated cells) to bring them together. Combat requires more prior knowledge as it uses the treatment information (e.g. compound identifier, dose,…) as covariates. Treatment information are available in a real-world scenario at the well level, including which wells are used as controls. We apply TVN on the embeddings (Ando *et al*., 2017) as follows. All embeddings of the untreated cells are whitened by computing the PCA basis so that these embeddings have zero mean and unit variance. The same affine transformation is then applied to all embeddings. Like in Tabak *et al*., (2019) but unlike Ando *et al*., (2017), we do not use CORAL (Sun *et al*., 2016) as we do not have enough embeddings per plate to compute a positive-semidefinite covariance metrics. Instead, for further correcting the batch effect, we are using Combat (Johnson *et al*., 2007). Combat adjusts the batch effect by using empirical Bayes methods. As covariates, we used treatment metadata (e.g. compound and concentration). Unlike other methods, we applied batch correction during training at each epoch before dimensionality reduction. The effect of the batch correction can be seen in **Supplementary Figure 4**.

#### 2.1.5 Dimensionality reduction of the embedding

To further improve clustering performance, we tested two dimensionality reduction methods, PCA, and t-SNE (Maaten and Hinton, 2008). We implemented an adaptive t-SNE that takes only the number of components that contain at least 95% of the variants and outputs three components.

### 2.2 Image data and preprocessing

#### 2.2.1 BBBC021 Image Dataset

We evaluated UMM Discovery on the BBBC021 (Caie *et al*., 2010) cellular dataset available from the Broad Bioimage Benchmark Collection presented in (Ljosa *et al*., 2013). The dataset consists of high content images from MCF-7 breast cancer cells exposed for 24h to a variety of chemical compounds (drugs). The cells are treated on 55 96-well plates across 10 batches. Three single-channel images are available representing fluorescence bound DNA, Tubulin and Actin. The total dataset contains 113 small molecules at eight concentrations resulting in 901 treatments representing 13200 field of view imaged over three channels. A subset of treatments in BBBC201 are annotated with their MOA. This subset contains 38 compounds dosed at various concentrations resulting in 103 treatments (3848 field of view), representing 12 known molecular MOA. For half of those, the MOA annotation was performed by visual inspection. The annotation of the other half was captured from literature. Ljosa *et al*., (2013), Ando *et al*., (2017), Godinez *et al*., (2017), and Tabak *et al*., (2019) also evaluated their methods on this gold standard subset.

#### 2.2.2 Preprocessing

We followed the approach undertaken by Ando *et al*., (2017) for preprocessing images. For each plate and each channel, a flatfield image is computed by taking the 10th percentile of all the intensity values and applying a Gaussian filter with sigma of 50. Each image is then divided by its respective flatfield image. Pixel intensity values lower than 1 are set to 1 before doing a log transformation in order to increase the dynamic range. To remove outliers’ values, any resulting values larger than 5 are clipped to 5. In addition to the preprocessing performed in Ando *et al*., (2017), the pixel intensities are image-wise and channel-wise normalized by z-score. Unlike Ando *et al*., (2017) and Tabak *et al*., (2019), no cell candidates were generated, which eliminates the time-consuming steps to locate and crop cells in the images. To reduce input size, images are tiled into 4 pieces. The advantage of tiling is that we can feed images with the highest resolution in the neural network without down-sampling them and no cell candidates have to be computed. The downside of tiling is that some tiles can hold almost no information. Aggregating embeddings for clustering analysis alleviate this problem.

### 2.3 Embedding Generation

After training the multi-scale net, all tiles are fed through the trained neural network to generate one representing embedding of size 64 for each tile. In case of the BBBC021, there are 4 tiles for each field-of-views and 4 field-of-view per well, resulting in 16 tile embeddings per well. As treatments are applied at a well level, we are mostly interested in well-level embeddings, and therefore we aggregated tile-level embeddings by computing their element-wise median, thus obtaining one embedding of size 64 for each well (**Fig. 2**). These well embeddings are used for clustering analysis to evaluate the performance of the clustering and to visually discover novel clusters. In this paper, we show that feature vectors can be used to identify MOAs of corresponding treatments.

**Fig. 2:**
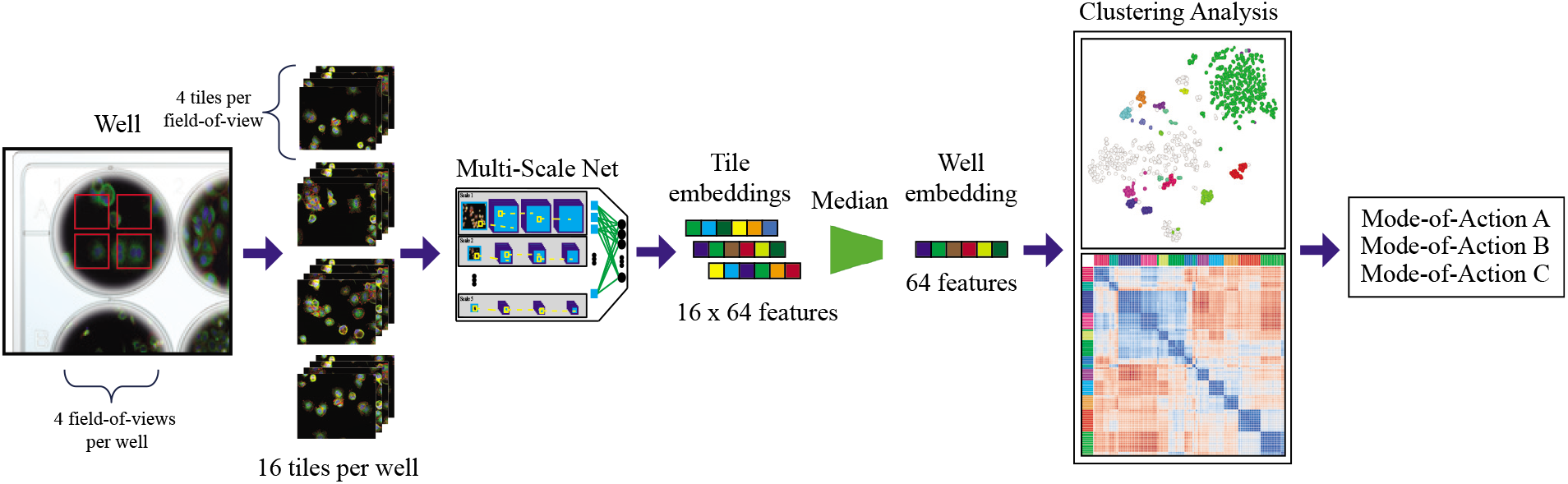
Embedding generation. Multiple fields-view are taken per well. These field-of-views are split into four tiles during preprocessing. All tiles of a well are forwarded through the neural network to create corresponding embeddings then aggregated to obtain one embedding per well. This well embedding is used for further analysis.

### 2.4 Evaluation Strategy

To evaluate the clustering quality, we made the assumption that images corresponding to the same MOA should cluster together, while images from different MOA should belong to different clusters. Suitable validation metrics take cohesion (intra-cluster) and separation (inter-cluster) into account (Palacio-Niño and Berzal, 2019). As each MOA represents one cluster, it is important to evaluate how close the embeddings of the treatments with the same MOA cluster together and separate from clusters with different MOA.

Another evaluation approach relies on using image metadata, by assuming that examples assigned to the same cluster have similar metadata (Palacio-Niño and Berzal, 2019). Because wells with the same treatment should cluster together, we computed the correlation between cluster assignment, MOA ground truth labels and treatment information (e.g. compound and dose).

#### 2.4.1 1-Nearest Neighbor Mode-of-Action Assignment

As in Tabak *et al*., (2019), Ando *et al*., (2017) and Ljosa *et al*., (2013), we used nearest neighbor classification to evaluate our method. Embeddings are first averaged at the plate level and then further aggregated by taking the median to obtain one embedding per treatment. We used the cosine distance to compute nearest neighbors. Then, to measure accuracy of MOA classification, we compared each example to its nearest neighbor, excluding examples treated with the same compound (NSC, <u>N</u>ot-<u>S</u>ame-<u>C</u>ompound) or examples either treated with the same compound or the same batch (NSCB, <u>N</u>ot-<u>S</u>ame-<u>C</u>ompound or <u>B</u>atch). MOA that are only present in one batch were excluded in the latter.

#### 2.4.2 Silhouette Score on Mode-of-Action

The silhouette score (Rousseeuw, 1987) evaluates how well the clusters are defined by using the mean intra-cluster distance *a*(*i*) and the mean nearest-cluster distance *b*(*i*) for sample *i* and is defined as:

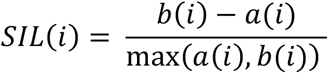

The range of the silhouette score is between −1 and 1. Negative values indicate that samples have been assigned to the wrong cluster. Positive values indicate that samples with the same assignment are clustering together and are well separated to samples with different assignments. As stated in Tabak *et al*., (2019), the silhouette score is more global than NSC/NSCB, as it takes global clustering into account instead of just the nearest neighbor of a sample. We used the silhouette score to evaluate the clustering at the well level. The well embeddings with the same MOA should cluster together and separate well from embeddings with different assignment.

#### 2.4.3 Completeness score on treatment metadata

The completeness score (Rosenberg and Hirschberg, 2007) is defined as

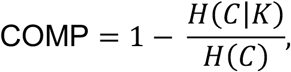

where *H*(*C|K*) is the conditional entropy of class *C* given cluster assignment *K* and *H*(*C*) is the entropy of class *C*. The completeness score on treatment metadata ranges from 0 to 1, where 1 indicates that all well embeddings with the same treatment are assigned to the same cluster. This score helps to evaluate if all embeddings with the treatment are clustering together independently of their batches.

#### 2.4.4 Adjusted Mutual information on compound metadata

The Adjusted Mutual Information (AMI) (Vinh *et al*., 2010) is based on information theory concepts and includes basic measures such as the entropy, the measure for disorder and the mutual information, and the measure of reduction in uncertainty. AMI is a modification of the mutual information as it ignores permutations and it is normalized against chance. It is defined as

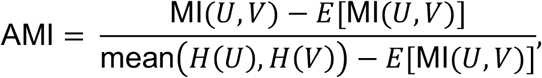

where U and V are the different cluster assignments, MI is the mutual information and E is the expected mutual information (Vinh *et al*., 2009). AMI allows us to measure the reduction in uncertainty about clustering assignments given the compound information from the metadata. The AMI score of a random assignment of clusters is close to zero, whereas similar clustering has a positive score and an exact match of cluster assignment and ground truth labels is 1. Therefore, the closer the AMI score is to 1, the better the pairwise correlation between clustering assignment and compound information is. In this case, one other beneficial characteristic of the AMI is that it penalizes over- and under-clustering. We used the AMI on the well embeddings and made the assumption that well embeddings of the same compound should cluster together or cluster close to DMSO when the treatment is not potent enough to have a visual effect of the MOA.

#### 2.4.5 Total clustering score

To identify how well the method performs with unlabeled data, we developed a new metric that we call total clustering score (TCS). The TCS consists of the adjusted mutual information, the silhouette score and the completeness and is defined as

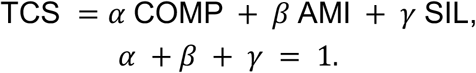

### 2.5 Best epoch determination

To avoid overfitting, we implemented a best epoch criterion. For the labeled dataset, the best epoch is selected by calculating the NSC for each epoch and taking the epoch with the highest score. For the unlabeled dataset this strategy cannot be applied. In this case, the TCS is used as best epoch criterion.

## 3 Results

First, we established a baseline model, which combines a multi-scale CNN (Godinez *et al*., 2017) with a K-means deep clustering backend (Caron *et al*., 2019) and applied it on the annotated subset of the publicly available cellular dataset BBBC021 (Caie *et al*., 2010). To evaluate the annotated clustering results, we used the NSC and SIL scores. After tuning the number of components for PCA and the number of seeds for K-means on the baseline model (**Supplementary Figure 2**), we evaluated the influence of different clustering algorithms (**Supplementary Table 2**).

Then, we incorporated batch effect removal. To asses that there is a batch effect and that it can be reduced with the combination of Combat and TVN, we performed a one-way ANOVA on the PCA components of the untreated well embeddings before and after batch correction with the previous mentioned methods. The **Supplementary Table 1** lists the F-value and the p-value of the analysis for the eight PCA components. All p-values before batch correction are significant (p<0.05), and therefore, we conclude that there are significant differences among batches. This is consistent with results from Ando *et al*., (2017) and Tabak *et al*., (2019). After batch correction, the F-values are considerably lower and the p-values (p>0.05) are not significant as we fail to reject the null hypothesis and conclude that after batch correction the untreated well embeddings have equal variances. In the **Supplementary Figure 3** can be seen that TVN and Combat decreases the batch effect. **Supplementary Table 3** shows the clustering results for TVN, Combat, and their combination (TVN followed by Combat and Combat followed by TVN). Every batch correction method significantly improves both NSC and SIL. Combat followed by TVN achieved the best results overall. Furthermore, we found that performing dimension reduction of the embeddings improved clustering performances (**Supplementary Table 4**).

In following section, we first show the results of the optimized UMM Discovery on the BBBC021 annotated subset, compare our method with other approaches and present how our method can be applied to unlabeled datasets for MOA discovery, where we applied it to the entire BBBC021 dataset.

### 3.1 BBBC021 annotated subset

We first evaluated UMM Discovery on the MOA annotated subset of the BBBC021 (Caie *et al*., 2010) image dataset. The best performances were achieved with using K-means for the clustering algorithm, adaptive t-SNE for dimensionality reduction and combining TVN and Combat for batch correction. In agreement with previous results (Caron *et al*., 2019), we found that allowing for a large number of seeds in K-means is necessary to achieve the best NSC. In our case, we set the number of seeds to the number of treatments. We achieved a NSC score of 97.09% and a silhouette score of 0.518 which indicates a close clustering of the same MOA and a clear separation between the clusters with different MOA. The COMP score was 0.936 and showed that well embeddings from the same treatments have the same cluster assignment. **Fig. 3** shows the NSC confusion matrix of this setup. UMM discovery was able to classify 100 of the 103 treatments’ MOA correctly to the right MOA. The t-SNE visualization of the well embeddings (**Fig. 4)** provides a visual confirmation of the clustering quality. The well embeddings of the same MOA are nicely clustering together and are clearly separated from the well embeddings of other MOAs.

**Fig. 3:**
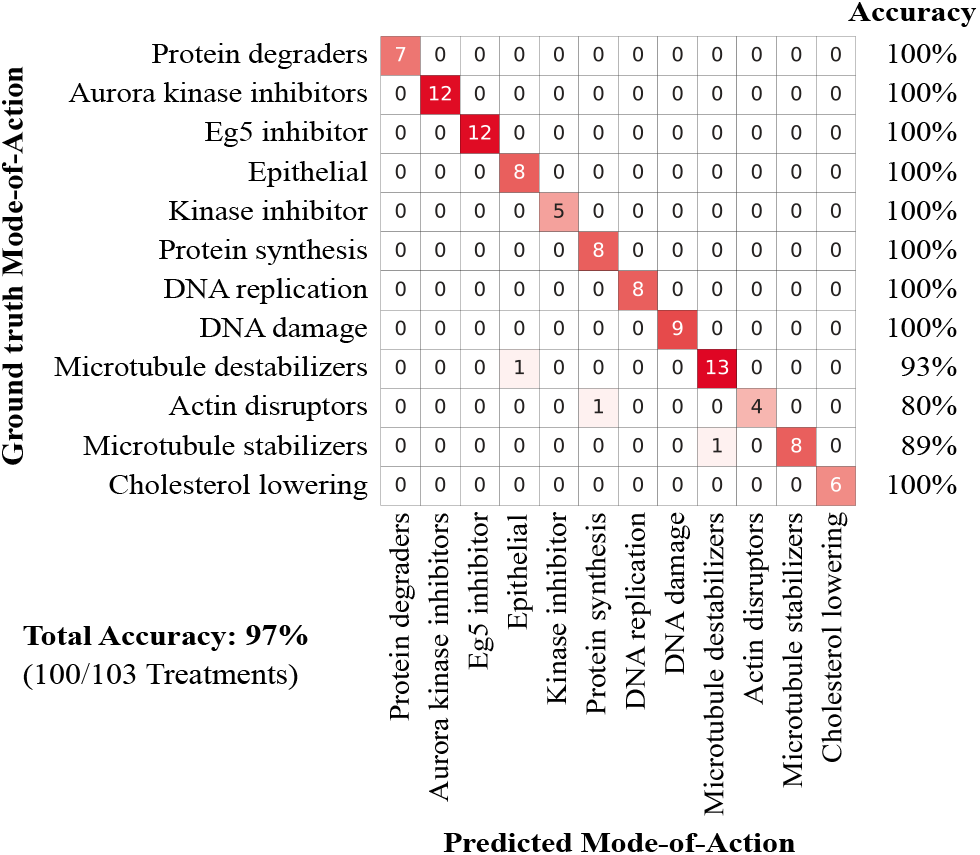
**NSC Confusion matrix** of the best performing setup. Almost all treatments are classified correctly by the first nearest neighbor NSC with the corresponding MOA label. Only three treatments were assigned to the wrong MOA.

**Fig. 4:**
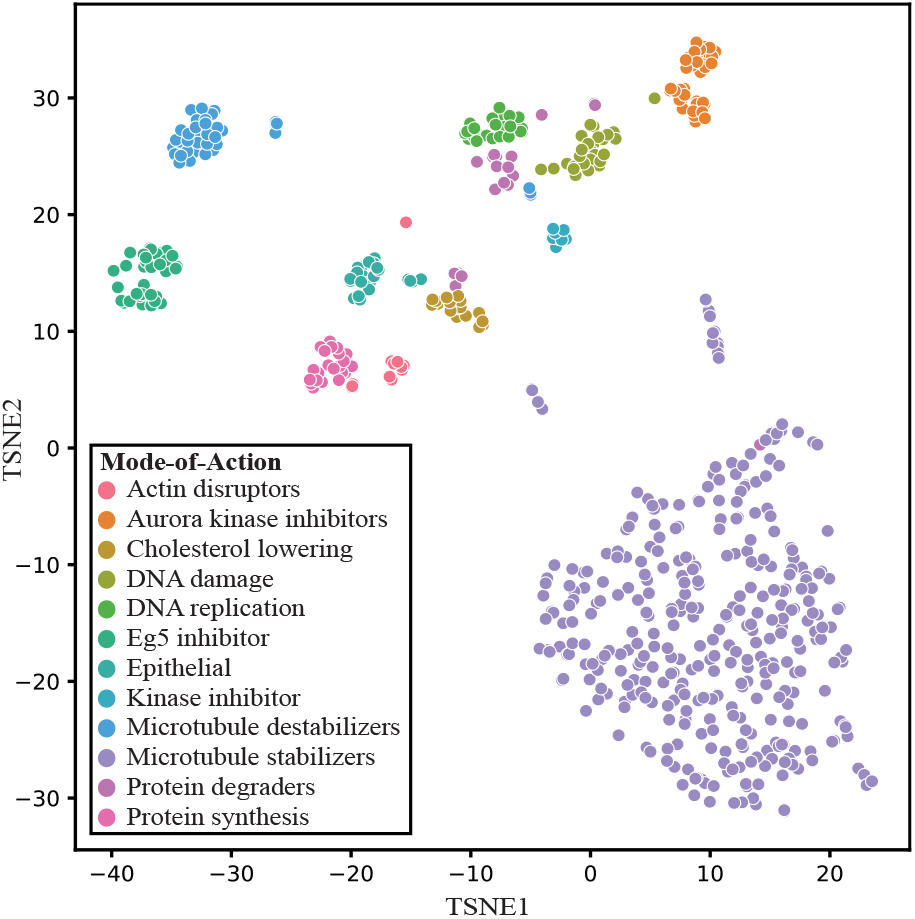
**t-SNE visualization of the well embeddings** of the annotated subset of BBBC021 without untreated cells of the best performing setup. Each dot corresponds to a well and is colored by MOA.

#### 3.1.1 Comparison with supervised training

We compared our method with a supervised training algorithm, where we trained the same adjusted Multi-Scale neural network with ground truth labels, once with the MOA as in Godinez *et al*., 2017), and once with the treatment metadata (compound + concentration) as in Godinez *et al*., (2018). After supervised training, we removed the last classification layer and created embeddings from all the tiles. Additionally, we used the same batch correction as in UMM discovery during training. **Table 1** recapitulates the performances of these approaches. UMM discovery is achieving results very close to supervised training on the MOA labels and outperforms the supervised approach with treatment labels.

**Table 1:**
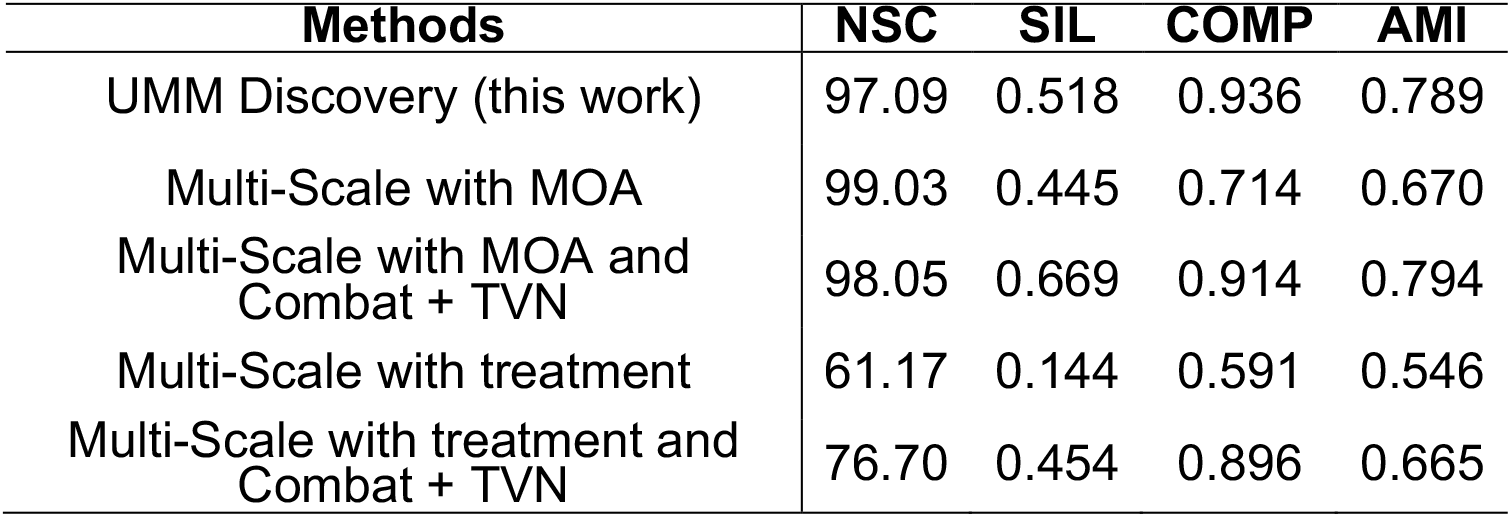
Comparing our work with supervised approach trained on the annotated BBBC021 subset with once the MOA as labels and once the treatment information as labels

#### 3.1.2 Comparison with other methods

We compared UMM Discovery with other state-of-the-art methods on the annotated subset of BBBC201. **Table 2** shows that UMM Discovery achieves the best performances in this comparison. One important distinction with other methods is that UMM Discovery is trained from scratch in a fully unsupervised manner. Ando *et al*., (2017) and Pawlowski *et al*., 2016) are using neural networks pretrained on other data to create their embeddings. The results show that our method performs very well on the small annotated subset of the BBBC021 dataset. However, the real benefit of an unsupervised approach is obtained when applying it to unlabeled datasets. Next, we evaluate UMM Discovery on the full BBBC201 dataset.

**Table 2:**
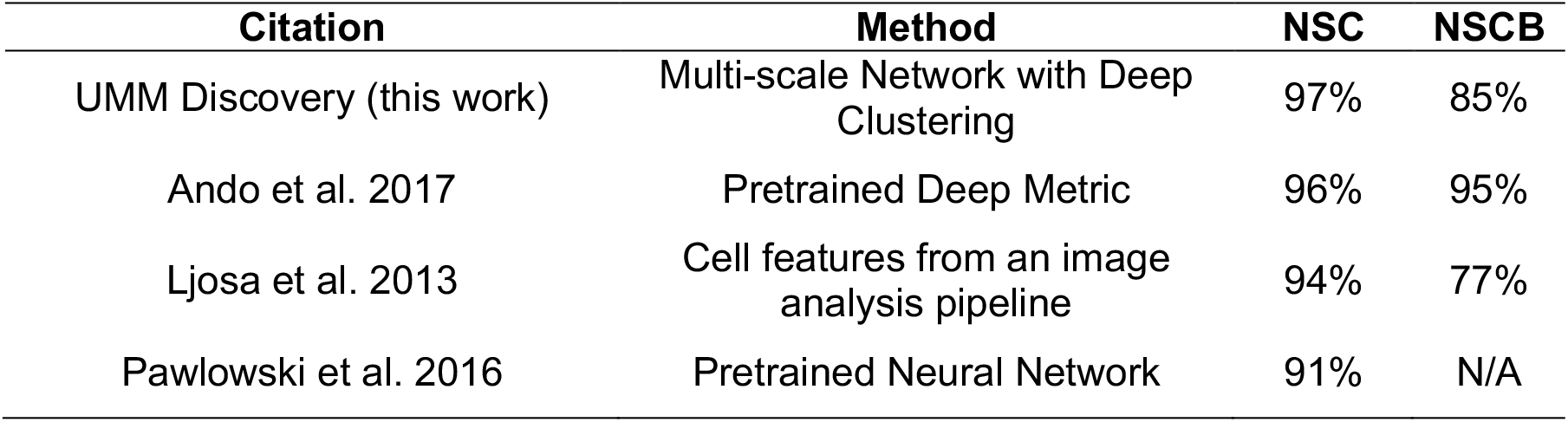
Comparison the results of UMM Discovery with other methods on the annotated subset of BBBC201.

| Citation | Method | NSC | NSCB |
| --- | --- | --- | --- |
| UMM Discovery (this work) | Multi-scale Network with Deep Clustering | 97% | 85% |
| Ando et al. 2017 | Pretrained Deep Metric | 96% | 95% |
| Ljosa et al. 2013 | Cell features from an image analysis pipeline | 94% | 77% |
| Pawlowski et al. 2016 | Pretrained Neural Network | 91% | N/A |

### 3.2 Unsupervised evaluation of the entire BBBC021 dataset

To evaluate whether UMM Discovery performed well on unlabeled datasets, we trained it on the entire BBBC021 dataset. As ground truth MOA labels are only available for a small subset of the images, we evaluated the clustering performance of our method using the available metadata (i.e. treatment metadata). We used the TCS to select the best epoch to avoid overfitting. The method achieved a silhouette score of 0.412, an AMI of 0.672 and a completeness score of 0.94. The high completeness score indicates that almost all embeddings with the same treatment are assigned to the same cluster.

We visualized the clustering of the well embeddings with t-SNE (**Fig. 5**) and could identify **i)** clusters that contained several well embeddings with known MOA, **ii)** outlier well embeddings with unexpected clustering behavior based on MOA annotation **iii)** as well as completely unannotated clusters (novel clusters). These novel clusters could represent new MOAs that are absent from the annotated subset. We focused in this evaluation on seven different handpicked groups of well embeddings, denoted with boxes in **Fig. 5**. The compounds and their concentrations within each box are listed in **Table 3** where we highlighted the concentrations that belong to the MOA annotations of the BBBC021 subset. **Supplementary Table 5** contains the compound target from DrugBank (Wishart *et al*., 2018).

**Fig. 5:**
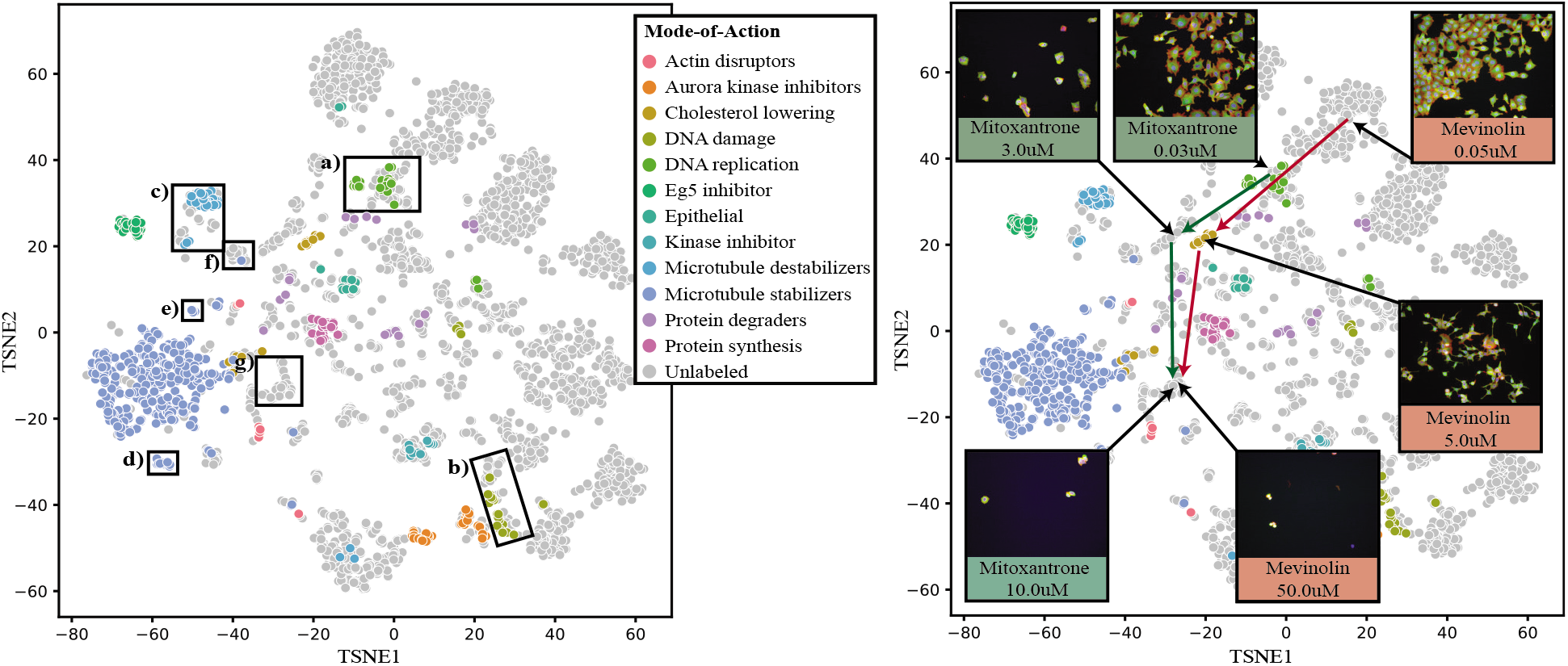
TSNE visualizations of the well embeddings of the entire BBBC021. Each dot corresponds to a well and is colored with the MOA labels of the BBBC021 subset. Grey dots correspond to the wells without MOA annotations. **Left:** The boxes indicate seven interesting clusters that we analyze in more detail. The compounds and their concentrations within each box are listed in **Table 3**. **Right:** Within the TSNE plot, a proliferation and a MOA effect can be discovered by tracking two compounds from the same batch that both induce fast cell death at high concentration. The arrows indicate the concentration path and the boxes show an example cellular image of the marked well embeddings for the corresponding compound and concentration.

**Table 3:** The compounds and the concentrations of the boxes in **Fig. 5**. The highlighted concentrations indicate well embeddings with annotated MOA ground truth from the BBBC021 subset. The colors of the highlighting are matching with the colors of the MOA in **Fig. 5**.

| | Compound | Concentration in $\mu\text{M}$ |
| --- | --- | --- |
| **a)** | camptothecin | 0.003, 0.01, 0.03 |
|  | mitoxantrone | 0.003, 0.01, 0.03, 0.1, 0.3 |
|  | floxuridine | 0.03, 0.1, 0.3, 1.0, 3.0, 10.0, 30.0, 100.0 |
|  | bleomycin | 0.15, 1.5, 5.0, 15.0, 50.0, |
|  | cathepsin inhibitor | 10.0 |
| **b)** | 5-flourouracil | 3.0, 10.0 |
|  | chlorambucil | 0.01, 1.0, 3.0, 10.0 |
|  | cisplatin | 0.03, 0.1, 0.3, 1.0, 3.0, 10.0 |
|  | etoposide | 0.3, 1.0, 3.0, 10.0 |
|  | mitomycin_C | 0.003, 0.01, 0.03, 0.1, 0.3, 1.0, 3.0 |
| **c)** | colchicine | 3.0 |
|  | demecolcine | 0.003, 0.01, 0.03, 0.1, 0.3, 1.0, 3.0, 10.0 |
|  | nocodazole | 0.03, 0.1, 0.3, 1.0, 3.0 |
|  | podophyllotoxin | 0.01 |
|  | taurocholate | 25.0 |
|  | trichostatin | 0.03 |
|  | vincristine | 0.003, 0.01, 0.03, 0.1, 0.3, 1.0, 3.0, 10.0 |
| **d)** | taxol | 0.3 |
| **e)** | taxol | 0.3 |
| **f)** | vinblastine | 0.003, 0.01, 0.03, 0.1, 0.3, 1.0, 3.0, 10.0 |
|  | taxol | 0.3 |
| **g)** | staurosporine | 0.03, 0.1, 0.3, 1.0 |
|  | tunicamycin | 50.0 |
|  | okadaic acid | 0.06, 0.2 |
|  | nystatin | 10.0 |
|  | mitoxantrone | 10.0 |
|  | mevinolin | 50.0 |
|  | filipin | 1.0, 3.0, 10.0 |

#### 3.2.1 MOA assignment of novel compounds

First, we tested if UMM Discovery is able to cluster unannotated compounds together with annotated compounds. Box (**a**) contains well embeddings that belong to three annotated compounds, namely *“camptothecin”, “mitoxantrone”* and *“floxuridine”*. The MOA label of all three annotated compounds is DNA replication. In addition to the annotated well embeddings, there are well embeddings of the same compound but with missing MOA annotation. The method was able to cluster the same compound with different concentrations that appear as same MOA together. It is worth noting that the batch correction is performed at the treatment level and therefore did not contribute to the co-clustering of these compounds. In the same cluster we can find well embeddings with unknown MOA, i.e., “*bleomycin*” and “*cathepsin inhibitor*”. If we look up the target of the unknown compound “*bleomycin*” in DrugBank, we find that it is “DNA ligase 1 inhibitor” and therefore fits perfectly in this cluster.

The second box (**b**) holds several well embeddings of the “DNA damage” compounds: *“cisplatin”, “chlorambucil”, “etoposide”* and *“mitomycin_c”*. Also, this box contains several unannotated concentrations from these compounds that our approach clusters nicely together. As unannotated compound, we found 5-flourouracil which has according to DrugBank the target “Thymidylate synthase”. This leads to DNA incorporation and destabilization and could reasonably be expected to induce the same phenotype as the other co-clustered compounds.

Furthermore, the unannotated compounds “*trichostatin*”, “*colchicine*”, *“taurocholate”* and *“podophyllotoxin”* are clustering together in box (**c**) with the known Microtubule destabilizer *“vincristine”, “nocadazole”* and *“demecolcine”*. The compounds *“colchicine”* and *“podophyllotoxin”* are both “Tubulin beta chain – inhibitors”, according to Drugbank (Wishart *et al*., 2018), which matches the assigned cluster. The full dataset contains 8 concentrations, 0.001-3mM, of the compound “*colchicine*”, where the concentration 0.03mM is annotated in the subset as Microtubule destabilizer. These annotated concentrations cluster close to the untreated wells. The clustering of the compound “*colchicine*” with a concentration of 3.0mM with other microtubule destabilizers and the deficiency of the annotated concentration 0.03mM in that cluster confirms the findings of Godinez *et al*., (2018). These three examples showcase that UMM Discovery is able to cluster unannotated compounds with annotated compounds, thus enabling potential MOA assignment.

#### 3.2.2 Identification of outliers

The compound “taxol” with the concentration of 0.3mM was used as positive control on each batch and has the MOA label Microtubule Stabilizer. Almost all well embeddings treated with “taxol” at 0.3mM are clustering together (big cluster colored in blue in **Fig. 5**). Box (**d**) indicates a sub cluster also containing well embeddings of the positive control. We compared the cellular images of this sub cluster with other cellular images of the positive control and discovered that they look very different (see **Supplementary Figure 5a**). Another outlier of the positive control is marked with box (**e**). The cellular images of this box are all blurrier (out of focus) in comparison with other cellular images of the positive control (see **Supplementary Figure 5b**). This result underscores the ability of the UMM Discovery to identify outliers.

#### 3.2.3 Discovery of a novel Mode-of-Action

Lastly, we evaluated the ability of UMM Discovery to identify novel phenotypes. Box (f) contains only the compound “vinblastine” at various concentration and one well embedding of the positive control (taxol at 0.3 mM) which is an outlier (see **Supplementary Figure 5c**). “vinblastine” is a breast cancer treatment that according to DrugBank (Wishart *et al*., 2018) inhibits the mitosis at the metaphase by binding the microtubular proteins of the mitotic spindle, which leads to crystallization of the microtubule and mitotic arrest. **Supplementary Figure 6** compares the cellular images of “vinblastine” with cellular images of the compound “vincristine” which is in a nearby clusters (see box **c**) and is annotated as Microtubule Destabilizer. The compounds “vinblastine” and “vincristine” have nearly identical molecular structure but shows a slightly different phenotype. This difference in phenotype is explained by the difference in drug affinities of these two compounds in DrugBank, whereby “vinblastine” was found to have a significant higher anti-tubulin activity than “vincristine” (Simmingsköld *et al*., 1981; Lobert *et al*., 1996).

The last box (**g**) is an exceptional case, as it consists of well embeddings from compounds in high concentrations. The target description from DrugBank (Wishart *et al*., 2018) for these compounds would not fit them together as same MOA. By looking at the images, we discovered that they contain almost no cells, suggesting strong toxicity and induced cell death from these compounds. As a result, the cluster appears to capture a strong proliferative effect rather than a MOA specific feature. The clustering of the compound *“staurosporine”* (at 0.1mM and 0.3mM) together with the compound *“mitoxantrone”* (at 10mM) was also detected in the work of Godinez *et al*., (2018). The right illustration of **Fig. 5** shows this proliferation effect in more detail. Displayed are two compounds, specifically “*mitoxantrone*” and “*mevinolin*”, with three different concentrations (high, intermediate, low) each. The embeddings of the low concentration lie close to each other, whereas the “*mitoxantrone*” in low concentration belongs to the “DNA replication” MOA. If we follow the concentration paths of these compounds indicated by the arrows, we find that the intermediate concentration is clustering in two different clusters. By looking at the cellular images of these well embeddings, we find that the cell density decreases from the low to the intermediate concentration and continues to shrink until there are almost no cells left in the high concentration cluster. This is the proliferation effect that our approach is able to capture thanks to the Multi-Scale Neural Network. Although the well embeddings with intermediate concentrations have similar cell density, the images can be clearly distinguished visually from each other and suggesting different MOA. The compound *“mevinolin”* at intermediate concentration is annotated as cholesterol lowering. The compound *“mitoxantrone”* at intermediate concentration clusters with the compound *“camptothecin”* at high concentration. This could fit as both of them have the same MOA, DNA replication, at lower concentrations (see box **a**). The results show that UMM Discovery is able to identify novel MOA and is able to distinguish between MOA effect and proliferation effect.

## 4 Discussion

In this work we present an unsupervised approach for the analysis of high-content cellular images based on the combination of two state-of-the-art methods, namely a Multi-Scale neural network architecture (Godinez et al., 2017) combined with a deep clustering framework (Caron *et al*., 2019). Our approach can accurately and robustly differentiate across a diverse set of phenotypes without the use of prior phenotypic annotation.

Unlike other approaches in the field, we propose a fully unsupervised method which starts from random initialization of the networks’ weights and learns to capture and distinguish the various drug-induced phenotypes from the intensity values of the cellular images alone. Further significant improvements were obtained by introducing batch correction methods which utilizes plate and compound annotation to reduce inter-plate effects. Yet at no stage is prior knowledge of MOA required or utilized.

We see two benefits from not using prior MOA annotation. Firstly, including MOA annotation in the training procedure runs the risk of reducing the space of discoverable MOAs to only those included in the training dataset. Secondly this inclusion is difficultly incorporated in a fashion enabling appropriate continuous interpretation of dose dependence. Indeed, drug concentration are typically either ignored (leading to MOA based clusters with little or no dose gradient) or treated as individual annotations included in the supervised training (leading to discrete dose dependent clusters with little overlap between iterative doses). In **Fig. 6** a TSNE visualization of the aggregated well embeddings of the BBBC021 subset is presented along various training epochs. Already in the initial untrained network (**Fig. 6a**) a clear separation between the negative control wells, in white, and some of the Mode-of-Actions, in the different colors, can be seen. As the training progresses (**Fig. 6a** to **Fig. 6d**), we can appreciate a very clear separation of the different mode of actions.

**Fig. 6.**
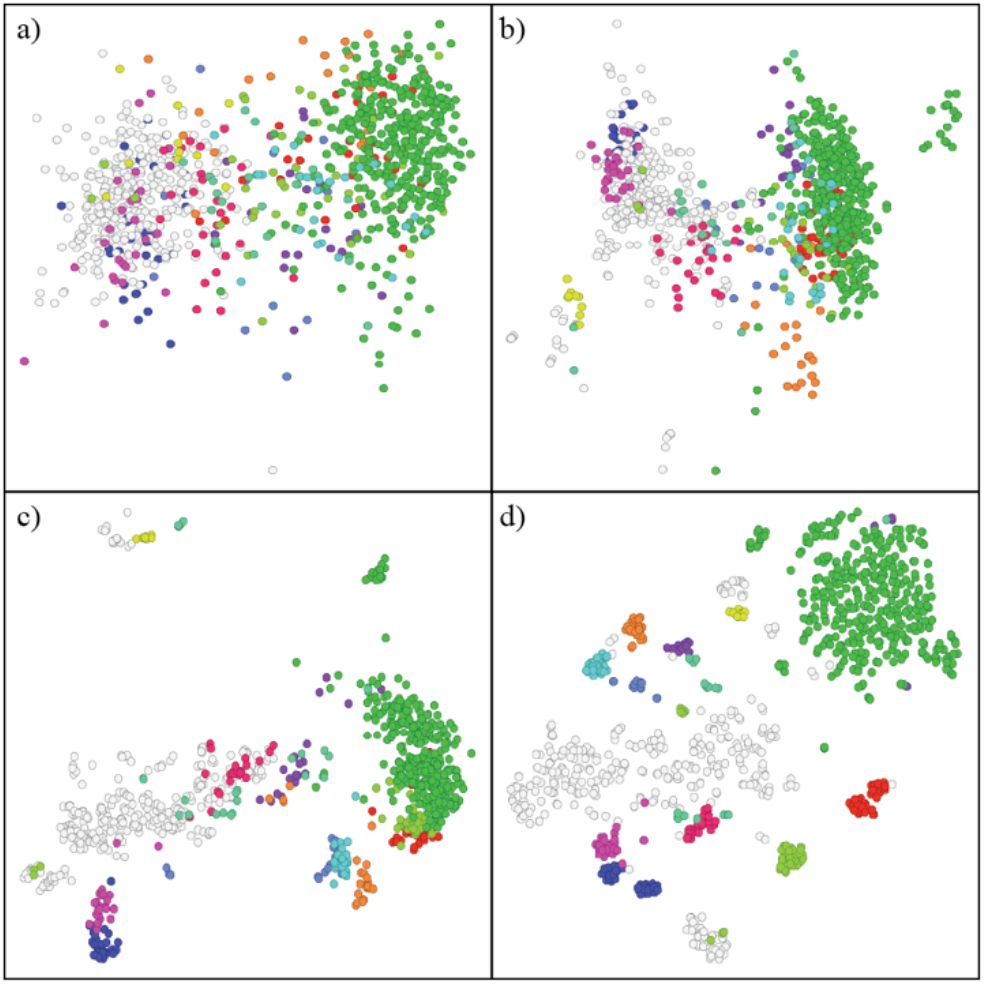
**t-SNE plot of the well embeddings during training,** where each point represents one well and is colored by the MOA annotation. White points correspond to untreated wells. a) Untrained neural network (initial weights), b) 1 epoch of training, c) 5 Epoch of training and d) after successful training.

Next the problem of batch effect was addressed. HCS datasets, like many high throughput biological datasets, are typically obtained from experiments with potential systematic and distinct noise introduced by technical heterogeneity (different experiment times, handlers, reagent lots, etc.). The BBBC021 dataset is no different and includes well annotated batches, which can be modeled using the differential effect observed across batches for the same treatment. In cases where such annotation would be lacking, considering individual plates as distinct batches would address the problem similarly. In our work performing such batch correction during training improved the results significantly. The batch correction not only increases the NSC for all clustering methods but also increases the silhouette score significantly, which implies that the overall clustering of the MOA is improved. Although, Combat or TVN alone could improve the NSC in all methods, the combination of both had a significantly higher silhouette score. Therefore, we continued our experiments with Combat plus TVN for batch correction. Additionally, it is important to mention that it is impossible to completely remove the batch effect on the dataset BBBC021 due to lack of sufficient coverage of identical compounds across batches. The application of a different batch correction methods, such as the implementation of the Wasserstein distance to correct for batch effect as in Tabak *et al*., (2019), could further improve the results. Regarding the dimensionality reduction, we tested PCA, t-SNE and UMAP, with t-SNE performing best. This would indicate the importance of local structure conservation rather than global structure to achieve higher performance. The relative underperformance of UMAP compared to t-SNE came as a surprise in this context, as both methods share some important characteristics, in particular local structure conservation. Better results might have been achieved with an exhaustive search of the UMAP parameter space. This was however not performed. The final optimized method outperformed image analysis pipeline and transfer learned methods as it achieves a NSC of 97% on the subset of the BBBC021. This result is very close to training in a supervised manner with MOA as ground truth labels. One unique aspect of UMM Discovery is the ability to perform training on the full BBBC021. Trained on the full dataset, we could show that similar phenotypes are clustering together and that our method is not only able to capture the MOA but also the proliferation effect. Additionally, we show that it can be used for Assigning MOA to novel unknown compound, identifying outliers and the discovery of novel MOA.

During the development of our method we tried several different setups. Yet still the clustering algorithms and their parameters, the dimensionality reduction methods, the batch correction methods and the hyperparameters of the neural network together represent an immense parameter space. We would like to mention that we did not conduct an exhaustive hyperparameter optimization. The hyperparameter that were used in scope of this work can be found in **Supplementary Table 6.**

While our approach shows promising results, we recognize that there are some limitations. One of the biggest limitations is the training time, as for each epoch we first have to pass all the images through the neural network to create the pseudo-labels by clustering their embeddings, to start the training. We trained our methods on the NVIDIA Tesla P100 with 16GB of memory and the training duration of our method on the annotated subset with batch correction and adjusted t-SNE as dimensionality reduction was approximately 23 hours. With PCA instead of t-SNE the training took approximately 18 hours. To address this issue, we are consequently exploring the inclusion of early pretrained layers instead of the initial weights. This would ease the evaluation of our method on larger datasets such as the BBBC036, consisting of five channels and 1553 different compounds. We have developed a fully unsupervised deep learning method based on two state-of-the-arts methods to cluster cellular image with similar Mode-of-Action together using only the pixel intensity values for the images. The method outperformed other existing methods applied to the same dataset. It achieved an NSC of 97.09% on the BBBC021 subset, achieving near supervised-level accuracy. In this work, we show that by incorporating metadata information to correct the batch effect during training, the overall clustering can be improved significantly. The benefit of the method in comparison to supervised methods is that it is not forcing treatments together when they look very different, as its clusters images together only by looking at their intensity values.

## Acknowledgements

We would like to thank William J. Godinez, Christopher Ball and Mark Bray for fruitful discussions. We also acknowledge the BBBC021 dataset provided by Peter D. Caie via the Broad Bioimage Benchmark Collection.

## Funding

This work has been supported by the Novartis Institutes for Biomedical Research

### Conflict of Interest

During their involvement related to this reported work, all authors were employees and shareholders of Novartis.

## Notes

https://data.broadinstitute.org/bbbc/BBBC021/

