## Supplementary data for "Fully unsupervised deep mode of action learning for phenotyping high-content cellular images"

**Supplementary Table 1:** ANOVA of the batch

| | No Correction | | Batch Correction | |
|---|---|---|---|---|
| PC Comp | F | p | F | p |
| PC1 | 64.63 | 1.07e-66 | 0.88 | 0.548 |
| PC2 | 23.76 | 5.33e-31 | 1.10 | 0.365 |
| PC3 | 11.65 | 7.56e-16 | 1.87 | 0.055 |
| PC4 | 4.65 | 8.0e-6 | 0.90 | 0.529 |
| PC5 | 6.10 | 6.35e-08 | 1.62 | 0.108 |
| PC6 | 13.55 | 2.03e-18 | 0.28 | 0.980 |
| PC7 | 2.95 | 2.22e-3 | 2.54 | 0.008 |
| PC8 | 7.75 | 2.53e-10 | 1.80 | 0.067 |

**Supplementary Table 2:** The 1st nearest neighbor NSC accuracy and the Silhouette score for each clustering algorithm with the best parameters. The clustering algorithm PIC showed promising results, but the Silhouette score indicated that the overall clustering was still insufficient.

| Method | NSC | SIL |
|---|---|---|
| K-means 208 | 80.77 | 0.070 |
| Power Iteration Clustering | 92.31 | 0.189 |
| HDBScan | 87.50 | 0.173 |
| Adaptive K-Means | 81.73 | 0.178 |

**Supplementary Table 3:** Compares the 1st nearest neighbor NSC accuracy and the silhouette score of the different clustering methods with the various batch correction methods. The first column contains the various methods. The second column contains the results without batch correction. The third column contains the results with Typical Variation Normalization (TVN) and the fourth the results with Combat. The fifth and the sixth contain the combination of these to batch correction methods in different orders.

| | No Correction | | TVN | | Combat | | TVN + Combat | | Combat + TVN | |
|---|---|---|---|---|---|---|---|---|---|---|
| Method | NSC | SIL | NSC | SIL | NSC | SIL | NSC | SIL | NSC | SIL |
| K-Means 104 | 69.23 | -0.059 | 82.69 | 0.152 | 91.35 | 0.294 | 90.39 | 0.412 | 87.37 | 0.395 |
| K-Means 208 | 80.77 | 0.070 | 89.42 | 0.112 | 91.35 | 0.276 | 92.31 | 0.476 | 93.27 | 0.391 |
| HDBScan | 87.50 | 0.173 | 89.42 | 0.237 | 91.35 | 0.308 | 87.38 | 0.462 | 90.55 | 0.516 |
| Power Iterative Clustering | 92.31 | 0.189 | 91.35 | 0.185 | 94.23 | 0.363 | 90.39 | 0.426 | 92.31 | 0.463 |
| Adaptive K-Means | 81.73 | 0.178 | 82.69 | 0.109 | 93.27 | 0.229 | 84.62 | 0.38 | 88.35 | 0.437 |

**Supplementary Table 4:** Comparison of PCA with 16 components and T-SNE with 3 components as dimensionality reduction by the 1st nearest neighbor NSC accuracy and the silhouette score.

| | PCA with 16 components | | T-SNE with 3 components | |
|---|---|---|---|---|
| Methods | NSC | SIL | NSC | SIL |
| K-means 104 | 87.37 | 0.395 | **97.09** | **0.518** |
| K-means 208 | 93.27 | 0.391 | **97.09** | 0.428 |
| HDBScan | 90.55 | 0.516 | 94.23 | 0.519 |
| PIC | 92.31 | **0.463** | 94.23 | 0.444 |
| Adaptive K-Means | 88.35 | 0.437 | 92.23 | **0.546** |

**Supplementary Table 5:** List all the compounds and their concentrations from the market boxes in the main paper in section novel MOA discovery. The last column contains the compound targets (max 3) according to Drugbank separated by "|".

| | Compound | Concentration | Target according to drugback.ca |
|---|---|---|---|
| a) | camptothecin | 0.003, 0.01, 0.03 | DNA topoisomerase 1 |
| | mitoxantrone | 0.003, 0.01, 0.03, 0.1, 0.3 | DNA - intercalation \| DNA topoisomerase 2-alpha - inhibitor |
| | floxuridine | 0.03, 0.1, 0.3, 1.0, 3.0, 10.0, 30.0, 100.0 | Thymidylate synthase |
| | bleomycin | 0.15, 1.5, 5.0, 15.0, 50.0, | DNA ligase 1 - inhibitor \| DNA - cleavage |
| | Cathepsin inhibitor | 10.0 | - |
| b) | 5-flourouracil | 3.0, 10.0 | Thymidylate synthase \| DNA - incorporation into and destabilization \| A RNA - incorporation into and destabilization |
| | chlorambucil | 0.01, 1.0, 3.0, 10.0 | DNA - cross-linking/alkylation |
| | cisplatin | 0.03, 0.1, 0.3, 1.0, 3.0, 10.0 | DNA - cross-linking/alkylation |
| | etoposide | 0.3, 1.0, 3.0, 10.0 | DNA topoisomerase 2-alpha - inhibitor \| DNA topoisomerase 2-beta - inhibitor |
| | Mitomycin_C | 0.003, 0.01, 0.03, 0.1, 0.3, 1.0, 3.0 | DNA - cross-linking/alkylation |
| c) | colchicine | 3.0 | Tubulin beta chain - inhibitor |
| | demecolcine | 0.003, 0.01, 0.03, 0.1, 0.3, 1.0, 3.0, 10.0 | - |
| | nocodazole | 0.03, 0.1, 0.3, 1.0, 3.0 | Extracellular calcium-sensing receptor |
| | podophyllotoxin | 0.01 | DNA topoisomerase 2-alpha - inhibitor \| Tubulin beta chain - inhibitor \| Tubulin alpha-4A chain - inhibitor |
| | taurocholate | 25.0 | - |
| | trichostatin | 0.03 | Histone deacetylase 8 \| Acetoin utilization protein |
| | vincristine | 0.003, 0.01, 0.03, 0.1, 0.3, 1.0, 3.0, 10.0 | Tubulin beta chain - inhibitor \| Tubulin alpha-4A chain - inhibitor |
| d) | taxol | 0.3 | Tubulin beta-1 chain - inhibitor \| Apoptosis regulator Bcl-2 - inhibitor \| Microtubule-associated protein 4 |
| e) | taxol | 0.3 | Tubulin beta-1 chain - inhibitor \| Apoptosis regulator Bcl-2 - inhibitor \| Microtubule-associated protein 4 |
| f) | vinblastine | 0.003, 0.01, 0.03, 0.1, 0.3, 1.0, 3.0, 10.0 | Tubulin alpha-1A chain - aduct \| Tubulin beta chain - adduct \| Transcription factor AP-1 |
| | taxol | 0.3 | Tubulin beta-1 chain - inhibitor \| Apoptosis regulator Bcl-2 - inhibitor \| Microtubule-associated protein 4 |
| d) | staurosporine | 0.03, 0.1, 0.3, 1.0 | - |
| | tunicamycin | 50.0 | Tyrosine-protein kinase Lck \| Serine/threonine-protein kinase pim-1 \| MAP kinase-activated protein kinase 2 |
| | Okadaic_acid | 0.06, 0.2 | - |
| | nystatin | 10.0 | Ergosterol - binder |
| | mitoxantrone | 10.0 | DNA - intercalation \| DNA topoisomerase 2-alpha - inhibitor |
| | Mevinolin/ lovastatin | 50.0 | 3-hydroxy-3-methylglutaryl-coenzyme A reductase inhibitor \| Integrin alpha-L - inhibitor |
| | filipin | 1.0, 3.0, 10.0 | - |

**Supplementary Table 6:** Hyperparameter. The parameter listed below are all the parameters that where tested in scope of this work.

| Parameter | Value |
|---|---|
| **Hyperparameter:** | |
| Number of features | 64 |
| Learning rate | 0.05 |
| Weight decay power | -5 |
| Epochs | 200 |
| Momentum | 0.9 |
| Loss function | Cross-Entropy |
| Optimizer | SGD |
| Input size | 512 x 640 |
| PCA | 8, 16, 32 |
| TSNE | 3, 4 |
| UMAP | 3, 4, 16 |
| **Parameter Kmeans:** | |
| Number of clusters | 39, 104, 208, 416 |
| **Parameter PIC:** | |
| Sigma | 0.2, 0.5 |
| Number of nearest neighbors | 5, 10, 15, 25 |
| **Parameter HDBScan:** | |
| Min cluster size | 12, 15 |
| Min sample size | 5, 3 |
| Number of nearest neighbors | 25 |
| **Parameter adaptive Kmeans:** | |
| Number of clusters end | 39 |
| Number of clusters start | 208 |
| Number of epochs for decay | 150 |

**Supplementary Figure 1:** The architecture of the Multi-Scale Neural Network.

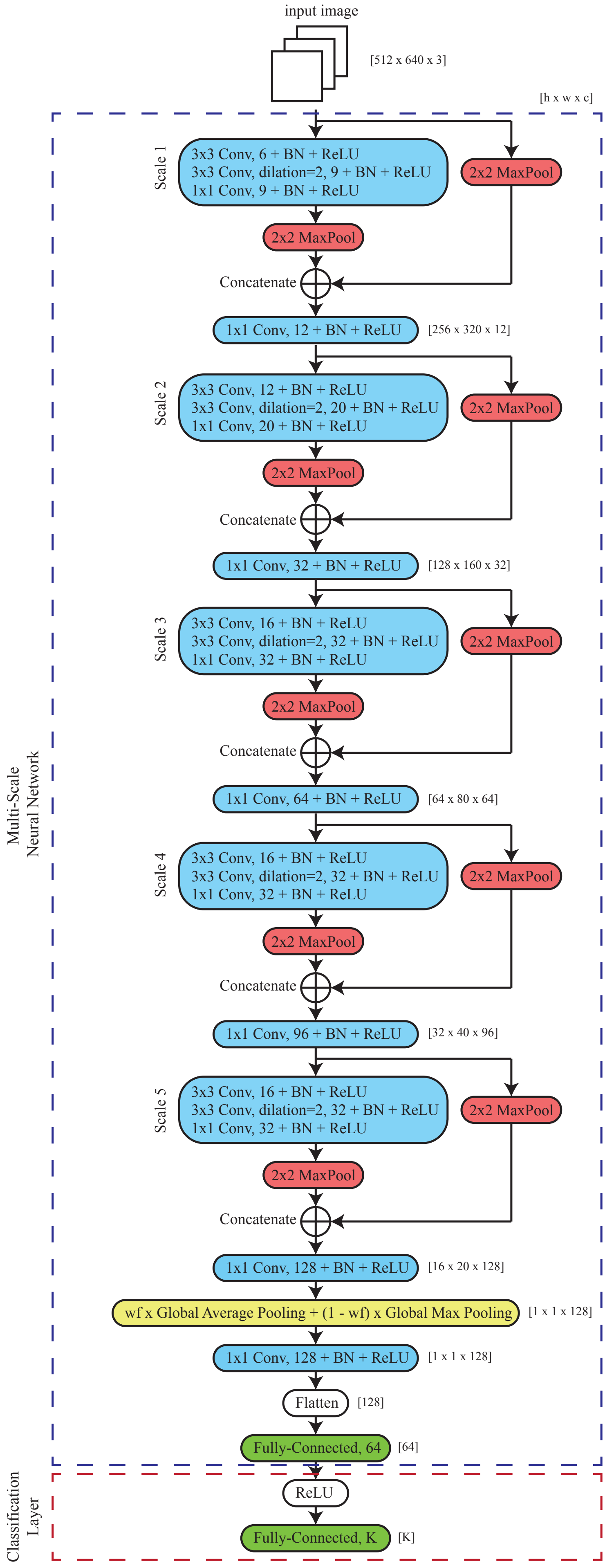

**Supplementary Figure 2: PCA and Seed determination**. The first nearest neighbor NSC accuracy by using a) different number of cluster seeds and b) Different number of principal components

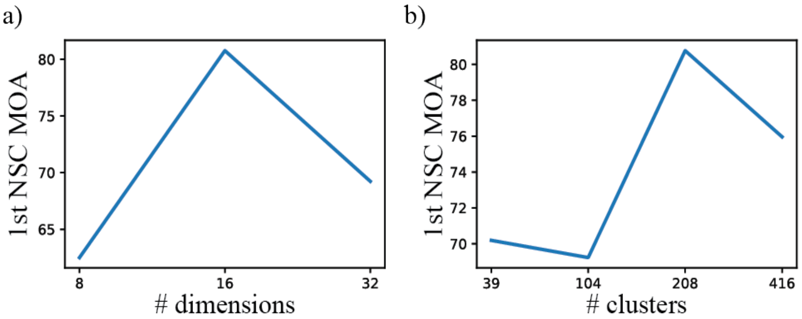

**Supplementary Figure 3: Batch effect removal within the untreated wells**. The PCA of the first and seconds components of the PCA of the untreated well embeddings after training the neural network. On the left a) No batch correction of the untreated well embeddings. There is variation between the different batches. On the right B) TVN and Combat together remove the difference in variances. The untreated well embeddings lie now on top of each other.

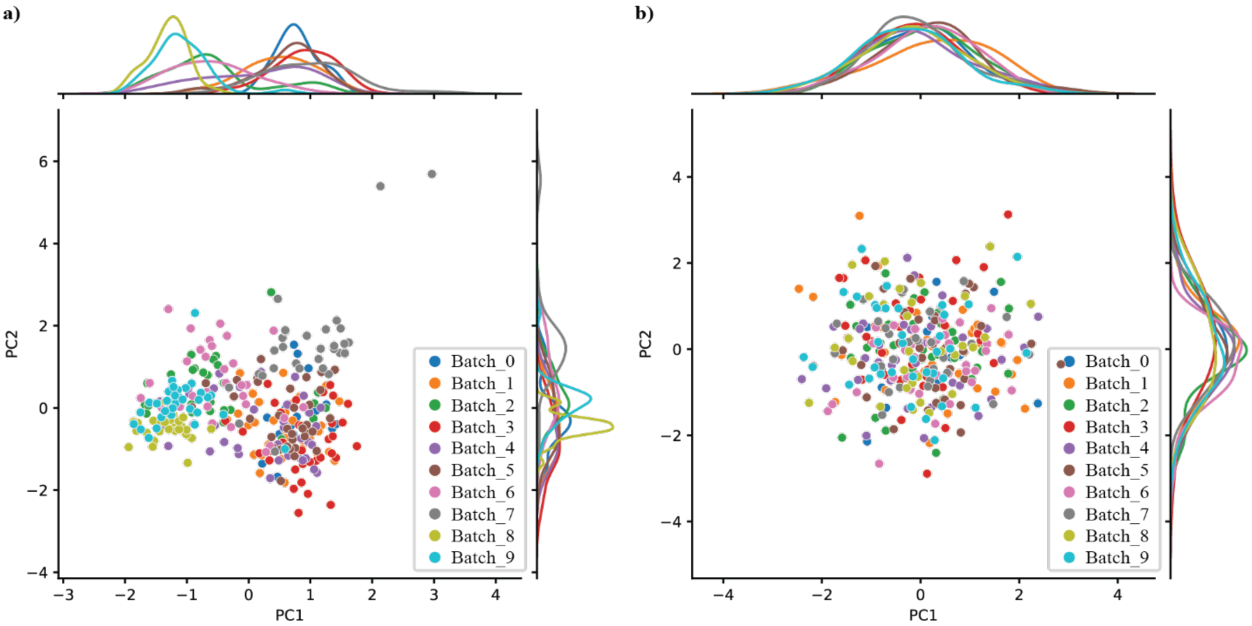

**Supplementary Figure 4:** Effect of batch correction illustrated on the untreated embeddings. Left: cosine distances between the untreated well embeddings. Right: cosine distance between the untreated well embeddings after TVN and Combat are applied.

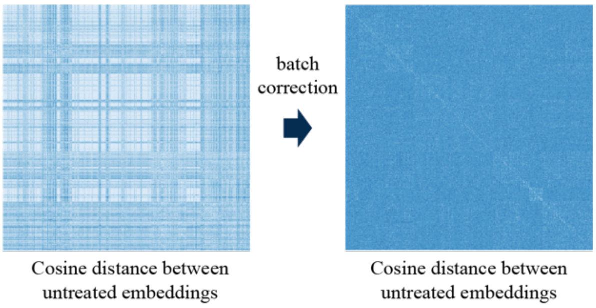

**Supplementary Figure 5: Comparison of cellular images of the positive control.** The top row shows typical cellular images of the positive control ("taxol" at 0.3uM) from different batches. The second row shows outliers of the positive control. a) very different MOA, b) out of focus and c) outlier close to well embeddings with the MOA Microtubule Destabilizers.

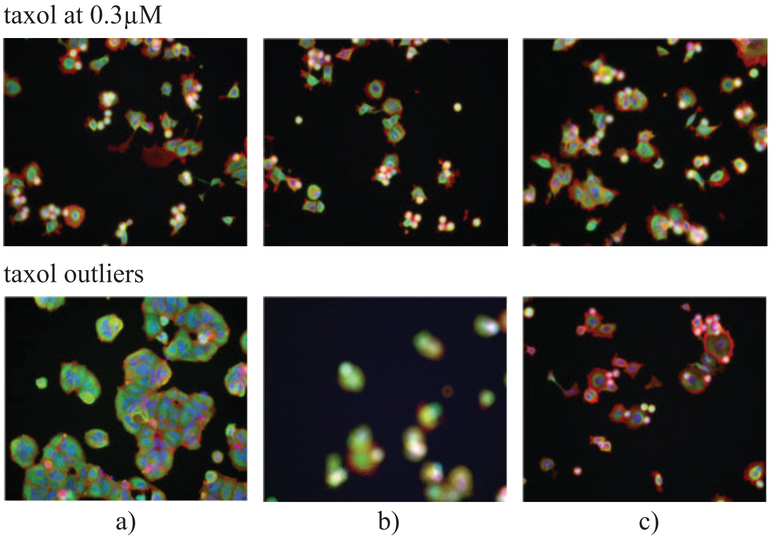

**Supplementary Figure 6:** Comparison of the compounds "vinblastine" and "vincristine" at different concentrations.

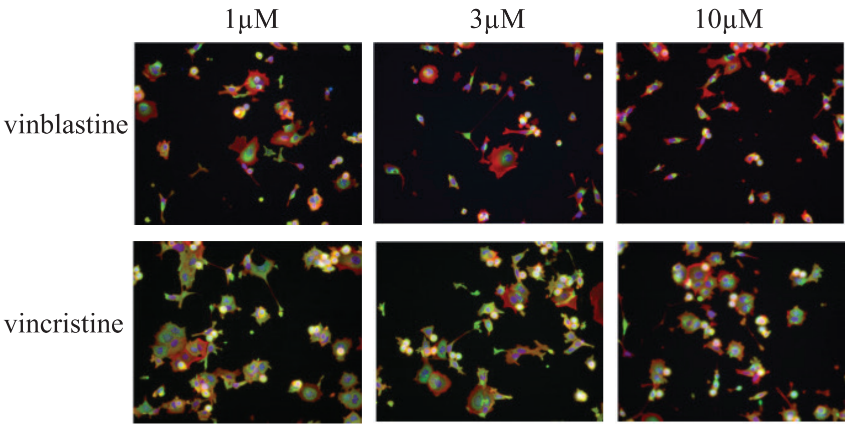
